# Experimental evolution under varying sex ratio and nutrient availability modulates male mating success in *Drosophila melanogaster*

**DOI:** 10.1101/2021.12.07.471570

**Authors:** Irem Sepil, Jennifer C. Perry, Alice Dore, Tracey Chapman, Stuart Wigby

## Abstract

Biased population sex ratios can alter optimal male mating strategies, and allocation to reproductive traits depends on nutrient availability. However, there is little information on how nutrition interacts with sex ratio to influence the evolution of pre-copulatory and post-copulatory traits separately. To address this omission, here we test how male mating success and reproductive investment evolve under varying sex ratios and adult diet in *Drosophila melanogaster* using an experimental evolution approach. We found that sex ratio and nutrient availability interacted to determine male pre-copulatory performance. Males from female-biased populations were slow to mate when they evolved on a protein-restricted diet. On the other hand, we found direct and non-interacting effects of sex ratio and nutrient availability on post-copulatory success, without interactions between them. Males that evolved on a protein-restricted diet were poor at suppressing female remating. Males that evolved under equal sex ratios fathered more offspring and were better at supressing female remating, relative to males from male-biased or female-biased populations. These results support the idea that sex ratios and nutrition interact to determine the evolution of pre-copulatory mating traits, but independently influence the evolution of post-copulatory traits.

## Introduction

The sociosexual environment has a profound impact on the strength and direction of sexual selection. Variation in the sex ratio alters the intensity of intra-sexual competition for mating opportunities and subsequent fertilization. Under male-biased (MB) sex ratios, heightened male-male competition is expected to select for male strategies that increase a male’s likelihood of securing a mating [1,2]. However, elevated polyandry – which typically occurs under MB sex ratios – can weaken pre-copulatory sexual selection on males, while strengthening post-copulatory selection [3]. In accordance with these findings, males are predicted to increase their investment in ejaculate under a MB sex ratio to ensure reproductive success in the presence of sperm competition [4–6]. Under a female-biased (FB) sex ratio, relaxed sexual selection is expected to result in reduced investment in competitive male adaptations for achieving high mating success and paternity.

Experimental evolution studies that manipulate the intensity of sexual selection are a powerful approach for investigating the evolution of reproductive strategies in response to varying sex ratios [7–10]. In agreement with theory, previous studies found that males from MB populations mated for longer, consistent with higher ejaculate investment [11,12]. Likewise, MB males depleted their ejaculates faster when mating multiply, suggesting elevated ejaculate investment in earlier matings at the expense of future reproductive performance [8,13]. Yet, in other studies, males from FB populations sired as many or more offspring than MB males, and MB males were slower to start mating compared with males from FB and equal-sex (EQ) populations [1,12,14].

These studies have shed light on evolved responses to the sociosexual environment. However, the evolution of male reproductive strategies is also likely to depend on the nutritional environment. In *Drosophila melanogaster*, males maintained on low-protein diets are poorer at securing matings [15]. Likewise, the impact of nutrient limitation on ejaculate investment is well established. A recent comparative study revealed that investment in seminal fluid protein production is highly sensitive to nutrient availability, whereas sperm quantity is only moderately affected [16]. Hence, the evolution of costly reproductive strategies is likely to be affected by the interaction between nutrient availability and the social environment [17]. However, this interaction has been rarely tested (but see [12,18]).

To examine how male reproductive strategies and investment in pre- and post-copulatory traits evolve under varying sexual selection and nutrient availability, we used experimental evolution in *D. melanogaster* [12]. Sexual selection and nutrient availability were manipulated by varying the adult sex ratio (MB, EQ or FB) and adult nutritional environment (a standard diet or a protein-restricted diet) in replicate populations [12]. We tested how male reproductive traits (mating success and latency, reproductive success, paternity, and suppression of female remating) diverged in response to sex ratio and adult nutrient availability. Because a male-biased sex ratio involves intense competition for few mating opportunities, we predicted that MB males should evolve to invest heavily in both securing matings (pre-copulatory reproductive traits) and ejaculate transfer when they do get a mating opportunity, and that they should have the highest reproductive output as a post-copulatory reproductive trait [19]. Alternatively, we predicted that elevated polyandry might weaken the selection on pre-copulatory traits but strengthen the selection on post-copulatory traits in MB males. We also predicted that an increase in investment would only be possible when nutrients are readily available, so the effects of sex ratio should be limited to populations evolving on a standard-protein diet, especially for traits affected by seminal fluid protein quantity or quality, such as remating latency [20,21].

## Material and Methods

### Experimental Evolution

Experimentally-evolving populations were derived from an outbred Dahomey (*Dah*) wild-type stock [12,18]. Populations evolved under one of three sex ratio treatments in combination with one of two adult dietary regimes in a fully factorial design. The adult diet treatments were either standard sugar-yeast-agar (SYA) medium or a protein-restricted SYA medium containing only 20% of the yeast content (detailed in [12,18]). Each population evolved at a male-biased (MB, 70 males:30 females), equal (EQ, 50:50) or female-biased (FB, 25:75) adult sex ratio. Each sex ratio and dietary combination had three replicates to give a total of 18 populations. To propagate each population, eggs were collected on the 8^th^ day after eclosion and larvae were raised at standard density on standard SYA [18]. Populations were assayed after 35 generations of experimental evolution. We used *sparkling* (*spa*) females and *spa* rival males for focal male mating and female remating assays. *spa* provides a recessive phenotypic marker for paternity assignment, and was backcrossed into the *Dah* background.

### Experimental Design

The main experiment was preceded by two generations in which all populations were reared under standardised conditions on Lewis medium supplemented with live yeast [22], to reduce variation from parental effects. After two generations, virgin focal (experimentally evolved) males from each population were collected at eclosion and housed in vials in single sex groups of 12 for 4-6 days. On the day before mating assays, we placed 6 to 8-day old virgin *spa* females in individual vials containing Lewis medium supplemented with live yeast. To begin mating assays, we added a focal male to each female vial and allowed flies 320 min to mate. We recorded mating success, mating latency, and mating duration. Mated *spa* females were allowed to lay eggs for two days within the same vial. We counted emerging offspring twelve days later to measure male reproductive output. After two days, we transferred each female into a new vial with two 8 to 10-days old *spa* virgin males and gave females a five-hour window in which to remate once. Remating occurrence, latency, and duration were then recorded. Remated females were allowed to lay eggs within the same vial for an additional two days and we counted and phenotyped the emerging offspring to measure paternity share. Matings and rematings were run over two days.

### Statistical Analyses

Data were analysed using R version 3.6.3 [23]. Mating latency, offspring number and remating latency were analysed using a generalised linear mixed effect model (GLMM) with a Poisson error distribution corrected for overdispersion. Mating duration and remating duration were analysed using a linear mixed effect model. Proportion of matings and proportion of rematings were analysed using a GLMM with a binomial error distribution. Paternity share was analysed using a GLMM with a binomial error distribution corrected for overdispersion. The initial model included diet, sex ratio and their interaction and day of the experiment as fixed effects, and replicate population as a random effect. All GLMMs (except binary responses) were initially checked for overdispersion, and when present an observation level random effect was introduced to control for it [24]. In all analyses, model selection was performed by backward stepwise elimination; non-significant (p>0.05) variables were eliminated from the model to arrive at the minimal adequate model. Day and replicate population remained in the minimal models to control for this variation.

## Results

### Male mating latency and success

The interaction between adult sex ratio and diet significantly affected male mating latency (χ^2^_2_= 11.3; p= 0.003). MB males were slower to mate, consistent with previous findings [12], whereas EQ males were faster to mate, regardless of their diet (figure 1a). However, diet impacted mating latency in FB males, and flies that evolved on a standard diet were faster to mate. We also found a significant interaction between sex ratio and dietary manipulation for male mating probability (χ^2^_2_= 6; p= 0.048). Among flies that evolved on a standard diet, FB males were more successful than MB males at securing a mating, whereas for flies that evolved on a protein-restricted diet, the mating success of each group was similar (figure 1b). In contrast to previous findings [12], we found no effect of either sex ratio or diet on mating duration (sex ratio: χ^2^_2_= 0.2; p= 0.868; diet: χ^2^_1_= 0.3; p= 0.551; interaction: χ^2^_2_= 0.9; p= 0.632) (electronic supplementary material, figure S2).

**Figure 1:**
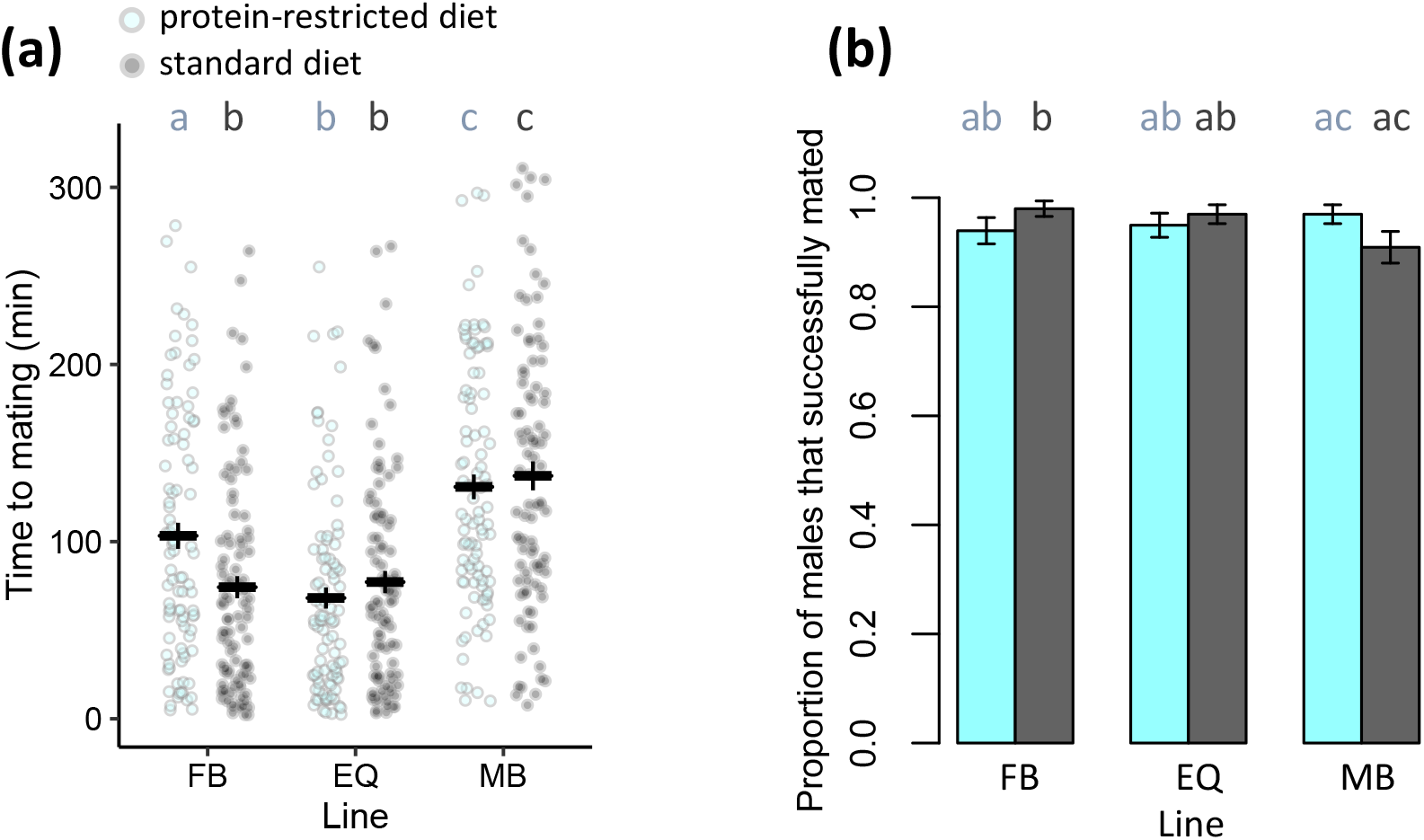
**(a)** Mating latencies and **(b)** mating success of experimentally evolved focal males (mean±se). Males evolved under female-biased (FB), equal (EQ) or male-biased (MB) sex ratios, and protein-restricted (20% yeast; light blue) or standard (100% yeast; dark grey) diet regimes. For both traits an interaction between sex-ratio and diet was detected. Letters indicate significant differences in post hoc tests. For plots split by replicate populations, see electronic supplementary material, figure S1.

### Male ability to induce female post-mating responses

Mates of EQ males produced more offspring, compared with mates of FB and MB males, in the 48h following a single mating (χ^2^_2_= 6.8; p= 0.033) (Figure 2a). Neither diet itself nor its interaction with sex ratio impacted offspring production (diet: χ^2^_1_= 0.5; p= 0.439; interaction: χ^2^_2_= 0.9; p= 0.608). Similarly, females that first mated with EQ males took longer to remate compared with females that first mated with FB or MB males (χ^2^_2_= 7.9; p= 0.018) (Figure 2b). Remating latency was not influenced by diet or its interaction with sex ratio (diet: χ^2^_1_= 0.1; p= 0.708; interaction: χ^2^_2_= 5.3; p= 0.068).

**Figure 2:**
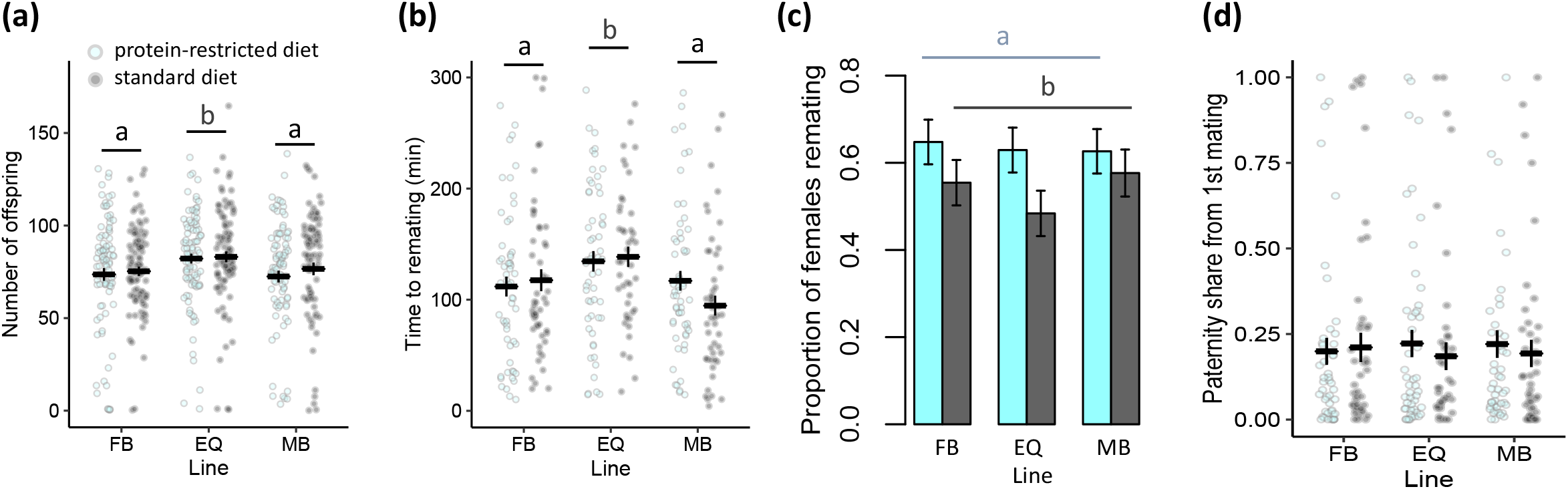
Female postmating response when wild-type females were first mated to experimentally-evolved focal males. **(a)** Number of offspring produced in 48 hours; **(b)** female remating latencies 48 hours following the initial mating; **(c)** remating probabilities 48 hours following the initial mating; **(d)** paternity share of the experimentally evolved focal males (mean±se). Males evolved under female-biased (FB), equal (EQ) or male-biased (MB) sex ratios, and protein-restricted (20% yeast; light blue) or standard (100% yeast; dark grey) diet regimes. Number of offspring and remating latencies varied as a response to sex ratio. Proportion of rematings varied as a response to adult diet. Sex-ratio and diet had no impact on paternity share when the focal males were the first to mate. Letters indicate significant differences among sex ratio (a, b) and diet treatments (c) in post hoc tests. For plots split by replicate populations, see electronic supplementary material, figure S3-S5.

However, we found that a female’s probability of remating was impacted by the focal male’s diet, such that females first mated to males that evolved on a protein-restricted diet were more likely to remate (χ^2^_1_= 5.3; p= 0.02) (Figure 2c). Remating probability was not affected by sex ratio or the interaction between sex ratio and diet (sex ratio: χ^2^_2_= 0.6; p= 0.728; interaction: χ^2^_2_= 0.8; p= 0.653). There was no significant effect of sex ratio, diet or their interaction on either paternity share (sex ratio: χ^2^_2_= 0.03; p= 0.982; diet: χ^2^_1_= 0.3; p= 0.571; interaction: χ^2^_2_= 0.8; p= 0.656) (Figure 2d) or remating duration (sex ratio: χ^2^_2_= 0.8; p= 0.655; diet: χ^2^_1_= 0.2; p= 0.593; interaction: χ^2^_2_= 2.1; p= 0.335) (electronic supplementary material, figure S6).

## Discussion

In contrast to our prediction, there was no evidence that MB males were better at securing a mating (indicating higher investment in pre-copulatory traits) or increased their post-copulatory investment. However, in line with the alternative prediction, we found that MB males were slower to start mating (indicating lower investment in pre-copulatory traits), and that these predicted effects only evolved when flies evolved on a standard-protein diet.

We also found evolved differences in male mating latency and success related to sex ratio, diet and their interaction. MB males were slower at achieving a mating, suggesting reduced pre-copulatory investment, consistent with recent findings of reduced pre-copulatory sexual selection in male-biased groups [3]. Likewise, our finding that males from FB populations were quicker to mate and more likely to mate than MB males (when they evolved on a standard-protein diet) is consistent with stronger pre-copulatory sexual selection on males in FB populations, possibly through selection on males to secure mating with the most fertile females [25]. Our results also suggest that evolving on a protein-restricted diet led to increased mating latency, consistent with the hypothesis that male pre-copulatory investment was limited by nutrient availability within FB populations.

To date, only few studies have used experimental evolution to simultaneously manipulate sex ratio and nutrition [12,18]. Here, we used the same experimentally evolving populations as these studies. In results congruent with ours, Rostant and colleagues [18] found that female resistance to male harm is more likely to evolve in standard-protein regimes, possibly due to the costs involved in expressing resistance. In our experiment, males that evolved on a standard-protein diet were better at suppressing female remating. Given that suppression of female remating is strongly driven by the transfer of seminal fluid proteins [26], populations that evolved on the protein-restricted diet are likely compromised in seminal fluid proteome production and transfer [27].

Other recent work on the same populations has revealed that MB males respond to the exposure of rivals by reducing their courtship behaviour and are hence slower to mate [12], congruent with our findings. This plastic response to the presence of rival males before mating can be explained by weakened pre-copulatory sexual selection with increased polyandry in MB populations [3], and by the frequent interruption of courtships by other males in these populations (see [12]).

Curiously, we found that EQ males were more successful with wild-type females than were MB and FB males in several reproductive traits, including offspring number and remating latency. Several hypotheses might explain this pattern. If wild-type females have an inverted U-shaped preference function for male reproductive traits, such that an intermediate optimum is favoured, then males from MB and FB populations that have evolved away from that optimum might suffer a disadvantage in courting and mating for wild-type females. Males from EQ populations that are more similar to wild-type populations (which have an approximately equal sex ratio [28]) might gain a relative advantage. An alternative hypothesis, that differences in effective population sizes among sex ratio treatments might limit the rate of evolution for some groups, is unlikely to explain the results because differences in effective population sizes among the treatments are relatively minor [29].

We found no difference in mating duration among experimentally-evolved populations. This is consistent with some previous experimental evolution studies [13,14], but inconsistent with two others, in which MB males mated for significantly longer than FB or EQ males [12,30]. Mating duration has been widely used as an indicator of ejaculate investment in *D. melanogaster;* however, it is becoming increasingly clear that it is not always a good proxy for sperm or seminal fluid protein transfer [19,31–33]. Although we found no differences in mating duration, the replicate populations differed in traits that suggest differential ejaculate investment, such as offspring number, remating latency and remating success, supporting the idea that there is no association between these measures and mating duration.

Despite the higher productivity and remating suppression of EQ males, we found that paternity share was similar among treatments. This result is consistent with previous work [14] and might be explained by the low heritability of traits determining sperm competition and the fact that sperm competition is still present within each sex ratio treatments [34].

In summary, we explored how consistent and long-term changes in the socio-sexual and nutritional environment interact to drive the evolution of male pre-copulatory and post-copulatory reproductive traits. Our results show that these environmental variables interact to drive the evolution of pre-copulatory traits such as mating latency and success, yet independently influence the evolution of post-copulatory traits, such as offspring number and remating latency.

## Authors’ contributions

The study was conceived and the experiments were conducted by I.S., J.C.P. and S.W.; the experimental evolution lines were supplied by A.D. and T.C.; the data was analysed by I.S. and A.D.; the article was written by I.S.; and edits on the drafts of the manuscript were provided by J.C.P., S.W. and T.C.

## Acknowledgements

We thank Wayne G Rostant and Janet S Mason for assistance in maintaining the sex ratio lines. We thank Eleanor Bath for assistance with data analyses.

