## Supplementary Figures 1-6 for "Experimental evolution under varying sex ratio and nutrient availability modulates male mating success in *Drosophila melanogaster*"

**(a)** **(b)**

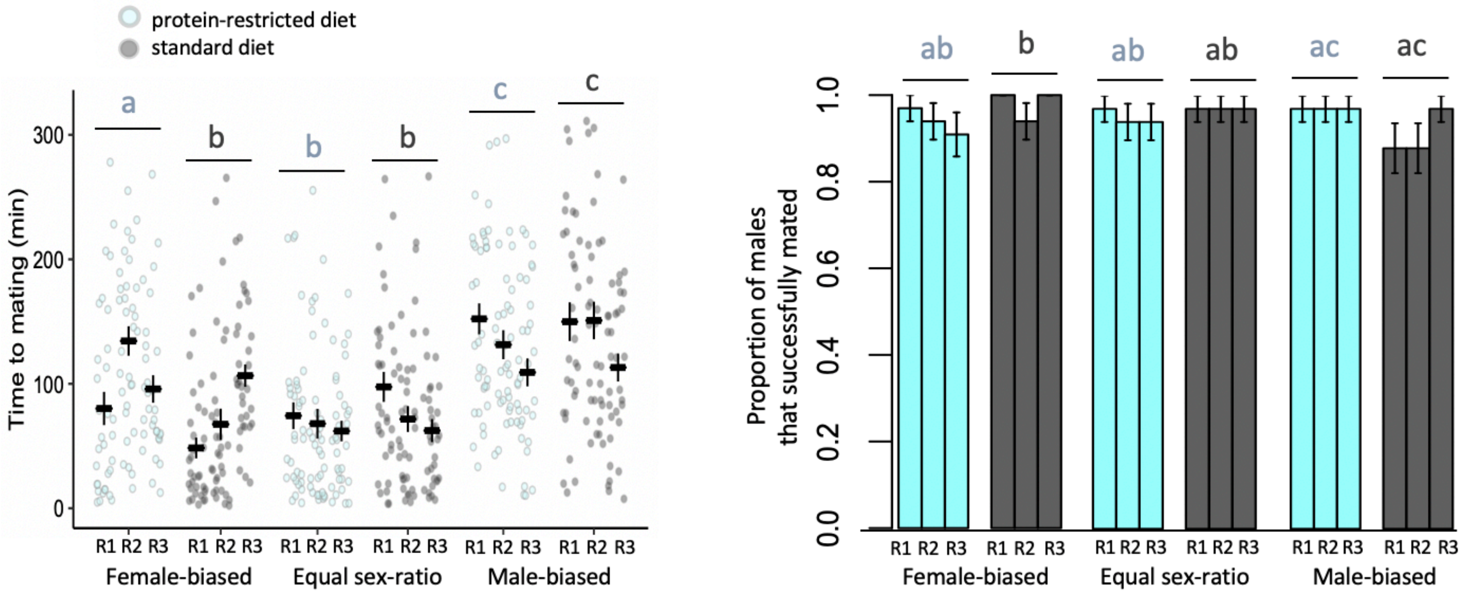


**Figure S1. (a)** Mating latencies and **(b)** mating success of experimentally evolved focal males from 18 populations differing in sex-ratios and adult dietary regimes (mean±se). Males evolved under female-biased, equal or male-biased sex ratios, and protein-restricted (20% yeast; light blue) or standard (100% yeast; dark grey) diet regimes. Each sex ratio and dietary combination had three replicates (R1-3). For both traits an interaction between sex-ratio and diet was detected. Letters indicate significant differences in post hoc tests.

**(a)** **(b)**

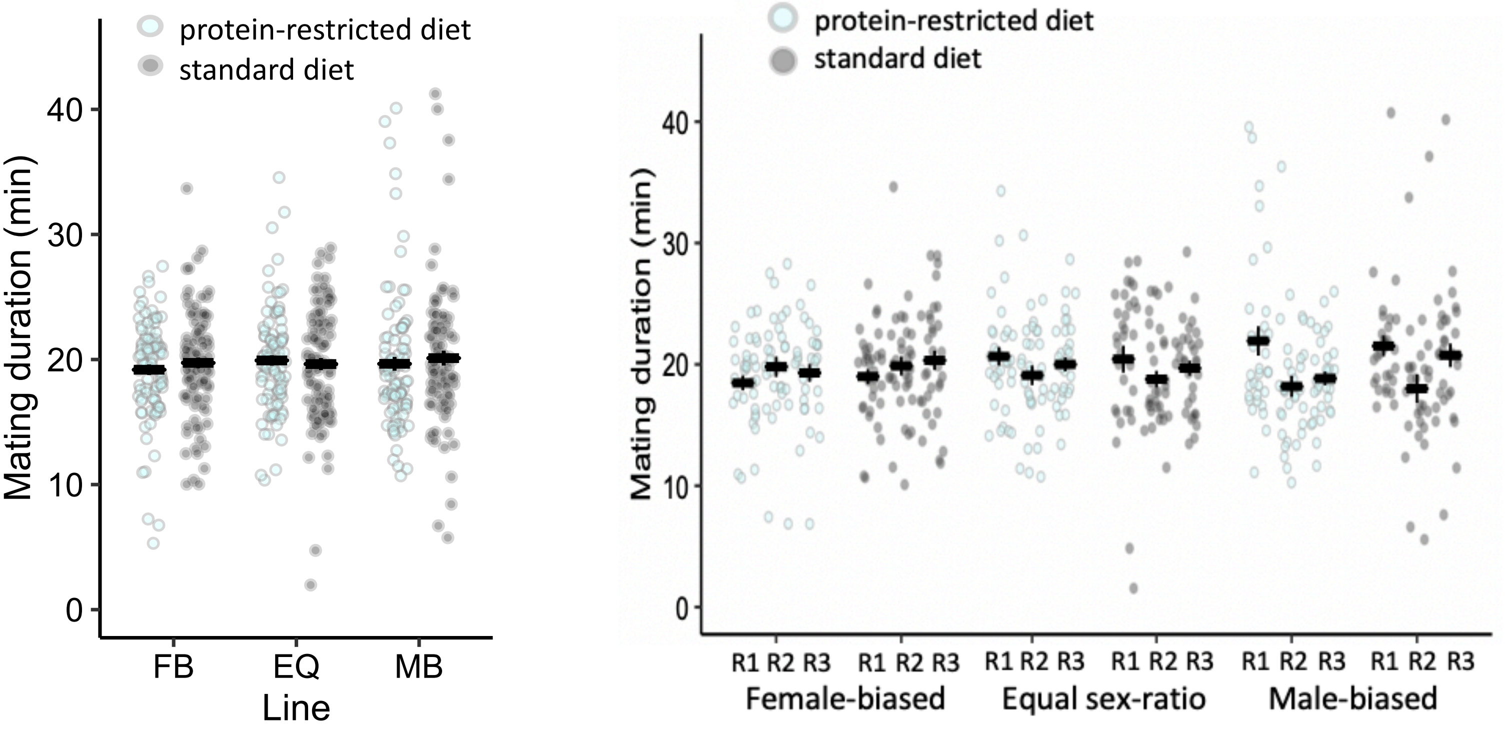


**Figure S2. (a)** Mating duration of experimentally evolved focal males (mean±se) **(b)** split by replicate populations. Males evolved under female-biased (FB), equal (EQ) or male-biased (MB) sex ratios, and protein-restricted (20% yeast; light blue) or standard (100% yeast; dark grey) diet regimes. Each sex ratio and dietary combination had three replicates (R1-3). Sex-ratio and diet had no impact on mating duration.

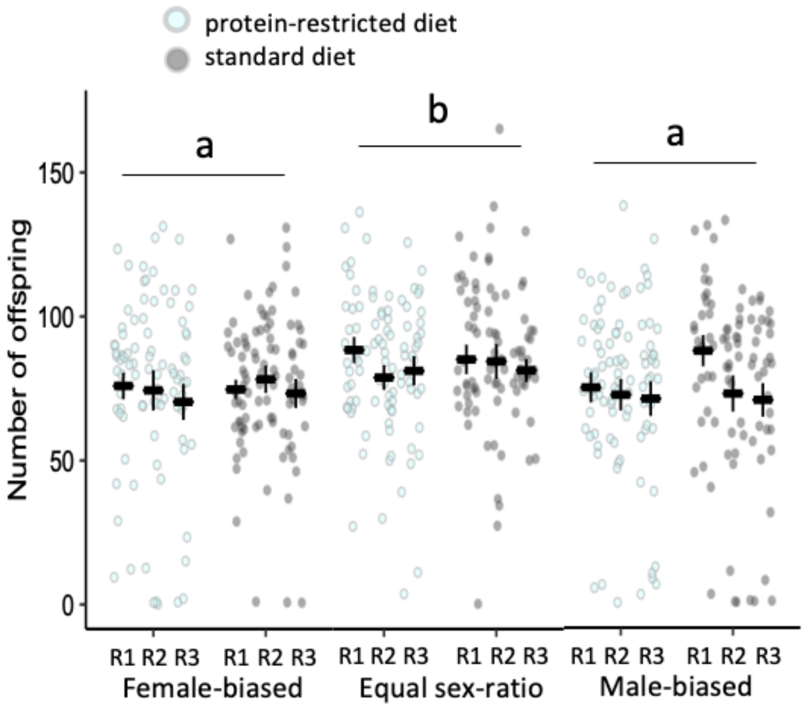


**Figure S3.** Number of offspring produced in 48 hours by experimentally evolved focal males from 18 populations differing in sex-ratios and adult dietary regimes (mean±se). Males evolved under female-biased, equal or male-biased sex ratios, and protein-restricted (20% yeast; light blue) or standard (100% yeast; dark grey) diet regimes. Each sex ratio and dietary combination had three replicates (R1-3). Number of offspring varied as a response to sex ratio. Letters indicate significant differences among sex ratio treatment in post hoc tests.

(a) (b)

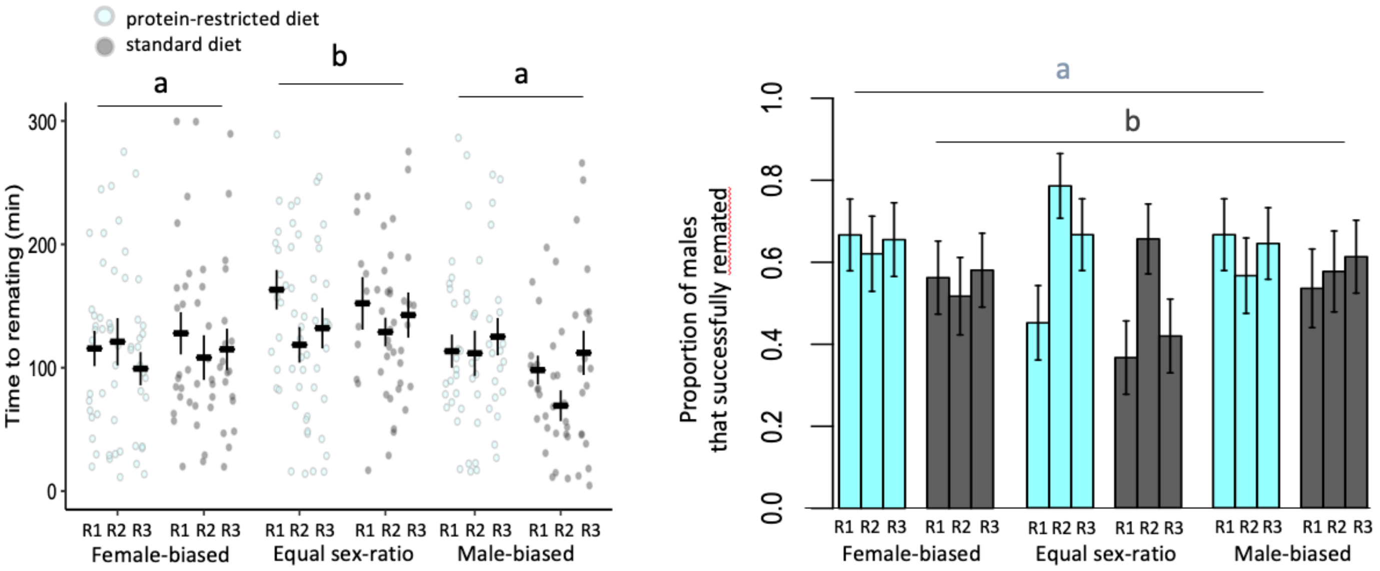


**Figure S4. (a)** Remating latencies and **(b)** remating success of experimentally evolved focal males from 18 populations differing in sex-ratios and adult dietary regimes (mean+se). Males evolved under female-biased, equal or male-biased sex ratios, and protein-restricted (20% yeast; light blue) or standard (100% yeast; dark grey) diet regimes. Each sex ratio and dietary combination had three replicates (R1-3). Remating latencies varied as a response to sex ratio. Proportion of rematings varied as a response to adult diet. Letters indicate significant differences among sex ratio (a) and diet treatments (b) in post hoc tests.

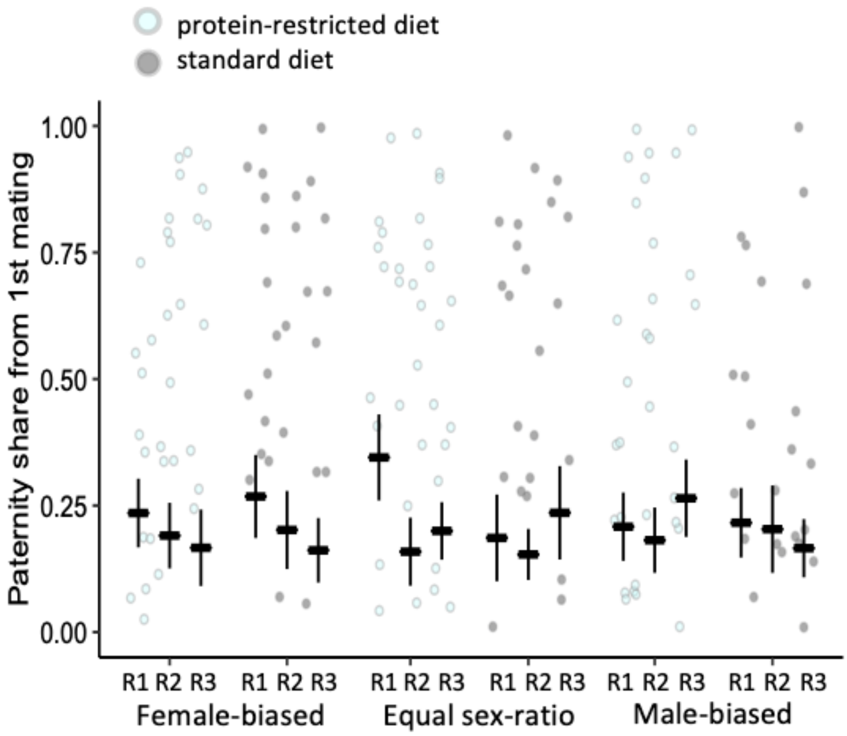


**Figure S5.** Paternity share of the experimentally evolved focal males (mean+se) from 18 populations differing in sex-ratios and adult dietary regimes (mean+se). Males evolved under female-biased, equal or male-biased sex ratios, and protein-restricted (20% yeast; light blue) or standard (100% yeast; dark grey) diet regimes. Each sex ratio and dietary combination had three replicates (R1-3). Sex-ratio and diet had no impact on paternity share when the focal males were the first to mate.

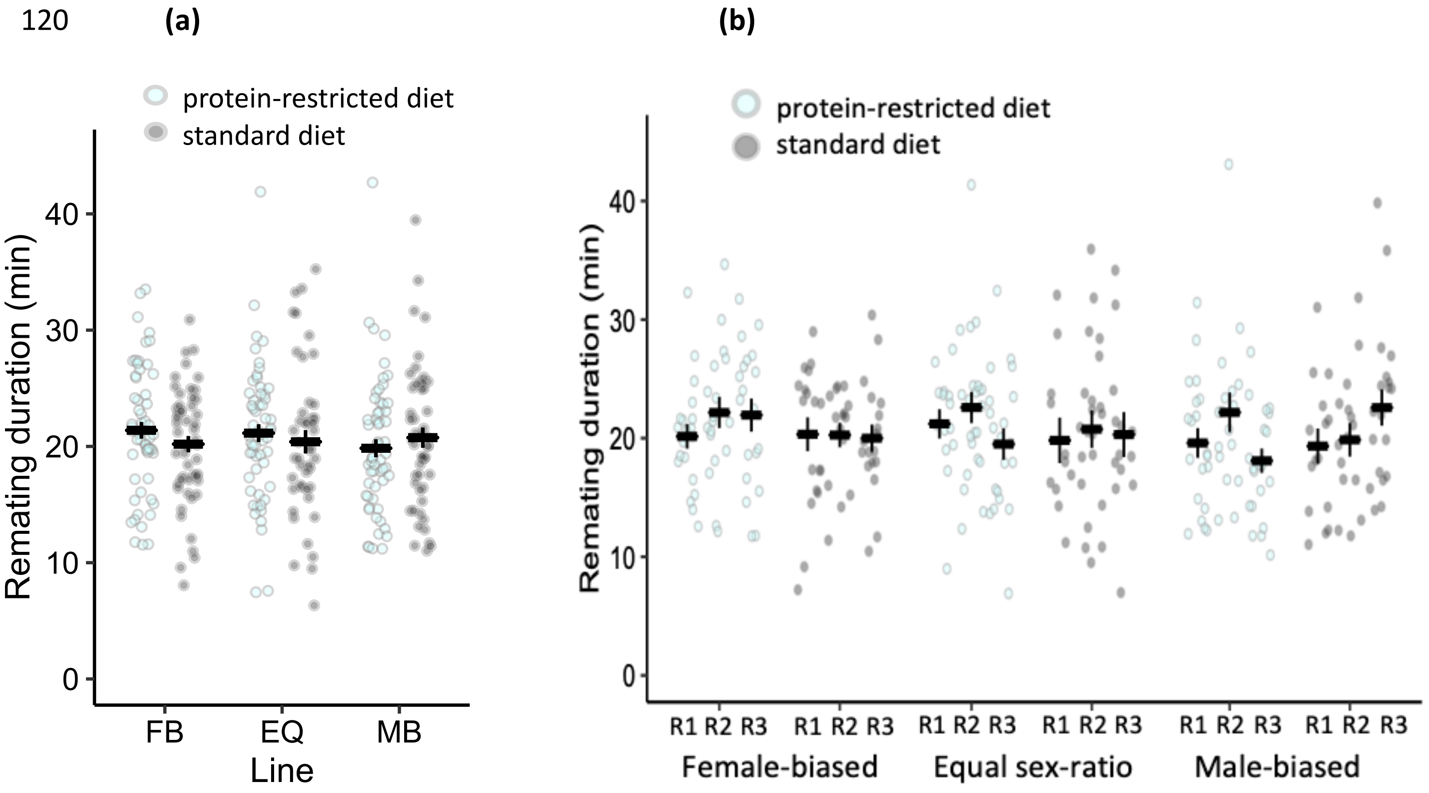


**Figure S6. (a)** Remating duration of experimentally evolved focal males (mean±se) **(b)** split by replicate populations. Males evolved under female-biased (FB), equal (EQ) or male-biased (MB) sex ratios, and protein-restricted (20% yeast; light blue) or standard (100% yeast; dark grey) diet regimes. Each sex ratio and dietary combination had three replicates (R1-3). Sex-ratio and diet had no impact on remating duration.
